# Changes in biophysical characteristics of YghA from *E. coli* due to variation in pH

**DOI:** 10.1101/2025.06.05.657471

**Authors:** Nadir Hasan, S. Bilal Jilani, Sana Saeed, Matthew R. Stoner, Syed Shams Yazdani, Yoonjoo Choi, Rizwan H. Khan

**Affiliations:** Interdisciplinary Biotechnology Center, Aligarh Muslim University, Aligarh, INDIA; International Center for Genetic Engineering and Biotechnology, New Delhi, INDIA; Combinatorial Tumor Immunotherapy MRC, Chonnam National University Medical School, Hwasun, SOUTH KOREA; Thayer School of Engineering, Dartmouth College, Hanover, New Hampshire, USA

**Author notes:** Equal contribution (Nadir Hasan, S. Bilal Jilani).

**Keywords:** Lignocellulose, tolerance, furan, bioenergy, inhibitor, industrial utility

## Abstract

YghA from *E. coli* has been reported to be involved in conferring tolerance against furan aldehyde inhibitors which are commonly generated in thermo-acidic pretreatment of lignocellulosic biomass. In this study we report the biophysical characterization of YghA from *E. coli* as a function of variation in pH. Fluorescence intensity of YghA increased by 2.5-fold in acidic pH as compared to circumneutral pH. In presence of the hydrophilic 8-anilinonaphthalene-1-sulfonic acid (ANS) dye, a 68 – 79.4-fold increase in fluorescence intensity at acidic pH was observed as compared to circumneutral pH, while at basic pH the increase was only 1 to 2-fold. Secondary structure analysis by circular dichroism signal at 222 and 208 nm suggests that the secondary structure of YghA is primarily composed of alpha-helix at pH 7 and the secondary structure is abruptly lost at pH 3 and lower. In agreement with these observations, MD simulations predicted greater structural variations at low pH when compared to neutral or high pH. Interface energy calculations using computational docking protocols suggested that YghA forms relatively more stable complex with NADH. The in silico pulling assay results also show that NADH is more preferred compared to NADPH. Molecular dynamics simulations also indicate that YghA is structurally unstable at acidic pH with significant variations in the root mean square deviation values of the tetramer backbone. Residues GLU 84 at pH 7 together with PRO 24 and LEU 236 at pH 1 were identified as flexible residues and are promising target for future mutagenesis studies targeted towards improving structural stability of YghA.

## 1. Introduction

Lignocellulosic carbon presents an inexhaustible source of carbon which can be channeled towards synthesis of organic compounds having industrial relevance (1–3). However, before the lignocellulosic carbon can be utilized by biological catalysts it needs to be processed via thermo-acidic pretreatment processes to release trapped carbon from cellulose and hemicellulose fractions (4–7). The process also results in formation of organic acids, aldehydes and aromatic compounds which can act as inhibitor of microbial metabolism (8–10). These inhibitors need to be converted into lesser toxic or neutral compounds before microbial utilization of lignocellulosic carbon can take place. Furfural is a potent furan aldehyde inhibitor generated due to breakdown of pentose sugars and acts synergistically to increase toxicity of other inhibitors present in the medium (11–14). From an industrial perspective, detoxification of furfural from the medium is essential before microbial production of a biomolecule can take place (15). Catalytic function of enzymes involved in wide functional aspects of microbial metabolism i.e. DNA repair (16–18), redox perturbation (19–22), polyamine metabolism (23–25), macromolecular synthesis (26), central metabolism (20,27) and transcription factors (28,29) have been implicated in furan tolerance in different microbial hosts.

Oxidoreductases class of enzymes perform an important role in maintaining redox homeostasis in the microbial cell in presence of furfural. Enzymes belonging to the alcohol/aldehyde reductases have been described with activity towards furan aldehydes and the catalytic activity has been associated with furan tolerance in the microbial host. Higher cellular levels of ADH1 (EC 1.1.1.1) and, ADH6 (EC 1.1.1.2) in *Saccharomyces cerevisiae* allowed growth to reach OD_600_ ∼ 1.5. In case of *Candida tropicalis*, purified ADH1 (EC1.1.1.1) has demonstrated activity against furfural and higher expression levels of ADH1 being correlated with furfural tolerance. Purified NADPH-specific butanol dehydrogenase (BdhA) from thermophilic *Thermoanaerobacter pseudethanolicus* 39E has shown furan reducing activity (40) and overexpression of the same led to ∼ 28% increase in microbial biomass and ∼ 25% increase in ethanol titers as compared to the control in *Caldicellulorisuptor bescii*. PsJN from *Pseudomonas putida* is an oxidoreductase which can convert aldehyde into corresponding acids and high levels of the same in microbial cells led to ∼ 1.5-fold higher microbial biomass as compared to the control strain in presence of furfural as sole carbon and energy source. Purified PsJN protein has been demonstrated to have furan reduction capability and overexpression being associated with furan tolerance in microbial cells (41).

FucO (EC 1.1.1.77) is a NADH-dependent L-1,2-propanediol reductase involved in fucose metabolism and has been shown to have invitro activity towards furfural (22). *Invivo* overexpression of FucO has been reported to increase bioethanol yield to 90% of the theoretical maximum along with an increase in the furfural tolerance in ethanologenic *E. coli* (30). YqhD (EC 1.1.1.2), an enzyme with a high degree of structural similarity to FucO, has been described as a major glycoaldehyde reductase in *E. coli* (33,35,36) and is induced as part of glutathione-independent response to lipid peroxidation (34). DkgA (EC 1.1.1. 346) has methylglyoxal reductase (31) and beta-keto ester reductase (32) activity. *In vivo* deletion of both *dkgA* and *yqhD* leads to two-fold increase in MIC values of ethanologenic *E. coli* (19) in presence of furfural and produced ethanol at ∼78% of the maximum theoretical yield as compared to the control strain with functional *dkgA* and *yghD* genes. Both DkgA and YqhD possess activity against furfural and have low apparent *K_m_* values of 23 and 8 µM towards NADPH which retards microbial growth due to scarcity of NADPH for biosynthetic reactions. Thus, enzymes without such strict cofactor requirements would be advantageous.

In recent studies, a novel oxidoreductase protein - YghA - has been reported to oxidize furfural in the presence of either NADH or NADPH (42). Overexpression of YghA in *E. coli* led to an increase in the maximum theoretical yield of ethanol from 69% in control strain to 97% observed in YghA overexpressing strain in presence of 1 g/L furfural. Due to the furfural detoxification property of YghA, it can serve as a promising candidate for enzyme immobilization to clear both furan inhibitors from the hydrolysate medium. It can serve as a cost efficient strategy where the enzyme is recycled to neutralize furfural in an organism independent manner. While promising, widespread adoption of YghA is limited due to the lack of data on biophysical characterization. In this study, we investigated the structural stability of the protein under different pH regimes. The study assumes significance since the thermochemical pretreatment process results in decrease of the pH of the medium. In addition, we report a few observations from a structural perspective. Firstly, *E. coli* YghA is highly similar to the *Salmonella enterica* YghA protein both in terms of sequence and predicted structural homology. Secondly, the structural stability of YghA protein is relatively resilient to the basic pH as compared to acidic pH. Thirdly, in silico studies suggest that NADH is the preferred cofactor as compared to NADPH for YghA protein.

## 2. Materials and methods

### 2.1 Strain, media and buffers

*E. coli* strain TOP10 (genotype F^-^, *mcr*A Δ(*mrr*-*hsd*RMS-*mcr*BC) ф80*lac*ZΔM15 Δ*lac*X74 *deo*R *rec*A1 *ara*D139 Δ(*ara-leu*)7697 *gal*U *gal*K *rps*L *end*A1 *nup*G was used as a host system (Invitrogen). Heterologous expression of YghA was achieved by transforming TOP10 with pTrcHisA plasmid harboring 6XHis-tagged YghA protein (42). The plasmid was first purified from strain DH5α and then transformed in TOP10 for protein purification. All strains were cultured at 37°C in LB medium in presence of 100 mg/L ampicillin. The buffer used for each pH condition is as follows: Glycine-HCl for pH1, 2 and 3; Sodium acetate for pH 4 and 5; Sodium phosphate for pH 6; Tris-HCl for pH 7 and 8; Glycine-NaOH for pH 9 and 10.

### 2.2 Purification of YghA

*E. coli* strain TOP10 harboring pTrcHisA-*yghA* was cultured overnight. 1% of the overnight culture was used to start a secondary culture. At OD_600_∼0.4 the secondary culture was induced by 0.1 mM IPTG for 4 hours. The cells were pelleted at 4°C and stored overnight in -80°C. The pellet was suspended in lysis buffer (5 mM imidazole, 500 mM NaCl, 20 mM Tris-Cl pH=7.0, 1 mg/mL lysozyme and 1 mM PMSF) and the cells ruptured using microtip sonicator. The lysate was centrifuged at ∼9,000g for 1 h at 4°C. The supernatant was filtered through 0.45 µm filter and the filtrate passed through Ni^2+^-NTA slurry via gravity flow. The resin was then washed with buffer (30 mM imidazole, 500 mM NaCl and 20 mM Tris-Cl pH=7.0). The bound YghA was finally eluted by elution buffer (250 mM imidazole, 500 mM NaCl, 20 mM Tris-Cl pH=7.0). The eluate was finally dialyzed against 100 mM phosphate buffer (pH=7.0). The purity of YghA protein as a single band was checked via visualization on SDS gel. The concentration of purified protein was determined by bicinchoninic acid (BCA) protein assay kit (G-Biosciences, MO, USA).

### 2.3 UV Visible spectroscopy

Ultraviolet-visible (UV-Vis) absorption spectra of protein samples were recorded using a Shimadzu UV/Vis-1900 spectrophotometer to investigate the impact of varying pH (2-10) on their chromophore environments. All samples maintained a protein concentration of 0.35 mg/ml. Spectra were acquired within the wavelength range of 240-400 nm using a 1 cm path length cuvette. Each measurement was performed in triplicate, and the resulting data were averaged for improved reliability.

### 2.4 Fluorescence spectrophotometer

Intrinsic fluorescence spectra of a protein sample (0.35 mg/mL) at varying pH (2-10) were obtained using a Shimadzu RF-5301 spectrofluorometer to elucidate the influence of pH on protein tertiary conformation. The samples were excited at 280 nm, and the emission spectra were acquired within the 300-400 nm range. Both excitation and emission slit widths were maintained at 5 nm. Each pH condition was measured in triplicate, ensuring data reliability and robustness for subsequent analysis.

### 2.5 ANS Assay

The surface hydrophobicity of a protein (0.35 mg/mL) at varying pH values (2-10) was assessed using 8-anilinonaphthalene-1-sulfonic acid (ANS) as a extrinsic fluorescent probe. Protein samples were incubated with ANS in a 1:30 molar ratio for 30 minutes under dark conditions to allow proper binding. Fluorescence spectra were then acquired using a Shimadzu RF-5301 spectrofluorometer with excitation at 380 nm and emission scanned from 400 to 600 nm. Both excitation and emission slit widths were set at 5 nm. Triplicate measurements were performed for each pH condition to ensure data reliability.

### 2.6 CD spectroscopy

Circular dichroism (CD) measurements were conducted using a Jasco J-815 spectropolarimeter. Prior to data acquisition, the instrument underwent a meticulous purge with high-purity (grade 3) nitrogen gas. Far-UV CD spectra were acquired at a protein concentration of 0.35 mg/ml under varying pH conditions (2-10). The spectral range spanned 190-250 nm, utilizing a 1 mm path length cuvette and a scanning speed of 200 nm/min, balancing signal-to-noise ratio with spectral resolution. Each measurement was performed in triplicate.

### 2.7 Structure Preparation and Computational Docking

We built a monomer structure of E-coli YghA (UniProtKB ID: P0AG84) using ColabFold (43). To generate a tetrameric form of YghA, the first ranked structure was then superimposed onto the crystal structure of *Salmonella enterica* YghA tetramer (PDB ID: 3R3S), which was co-determined with NAD. Assuming that the NAD binding pocket is the same for NADH and NADPH binding, the region was used for computational docking. The molecules were docked using the Rosetta ligand docking protocol (44) and FitDock (45) with default parameter settings.

#### 2.7.1 Molecular dynamics simulations

Molecular dynamics and pulling simulations were performed using GROMACS 2023 (46) with the AMBER99SB-ILDN force field and spce solvent model at 310K temperature and 1bar pressure. The best docked conformations of NADH and NADPH were used for Pulling simulations at pH 1, 4, 7, 10 and 12 to check effect and behavior of both the molecules within the pocket of YghA at different pH.

Monomer of *E. coli* YghA was used for the pulling simulations. To simulate the protonation states of each ionizable group at input pHs, the monomer was first uploaded to the PDB2PQR webserver (47), using the PropKa (48) method, hydrogen atoms were added to titratable residues. The produced files (.pqr) were transformed back into PDB files and set up in Gromacs for pulling MD simulations. Each system’s complex coordinate files were created, and ligands topology files were included in the topology files of the system. The complexes were boxed-in and solvated with water (49). Charge neutralization of the systems was achieved by adding NA and CL ions to the SOL (solvent) groups. The simulation systems were temperature-equilibrated at 310K using a modified V-rescale thermostat, pressure-equilibrated at 1 bar using C-rescale coupling, and energy-minimized using a steepest descent technique. During the phases of temperature and pressure equilibration (NVT and NPT), hydrogen bonding was restricted. Groups A and B were chosen for the pulling simulations; Group B represented the position-restrained YghA, and Group A represented the NADH or NADPH. Since we aimed to pull it out from the YghA pocket, Group A was not subjected to positional restraints. Pulling simulations were performed for 0.8 ns.

For MD simulations for the stability of tetramer structure at different pH levels, three systems were simulated for pH:1, 7 and 12.Equilibrations (NPT and NVT) were done at 310K temperature using V-rescale thermostat, and pressure at 1bar using C-rescale coupling after minimization of the systems using a steepest descent technique. After the simulation boxes were fully equilibrated, production MD was run for 100ns.

## 3. Results and discussion

### 3.1 Purification of YghA

YghA was purified using affinity chromatography with the elution profile as in Figure 1. Imidazole has potential to interfere with the absorption signal in the UV region and was removed by dialysis. To remove imidazole from our eluted fractions we pooled the three eluted fractions together and dialyzed against 100 mM sodium phosphate buffer (pH=7.2). To ensure complete removal of imidazole we switched the 1L dialysis solution 3-times in a 20h period. Purified YghA was checked after dialysis in SDS-PAGE to confirm its purity (>95% pure). Aliquots of the protein were snap frozen in liquid nitrogen and then stored in -20 till further analysis.

**Figure 1.**
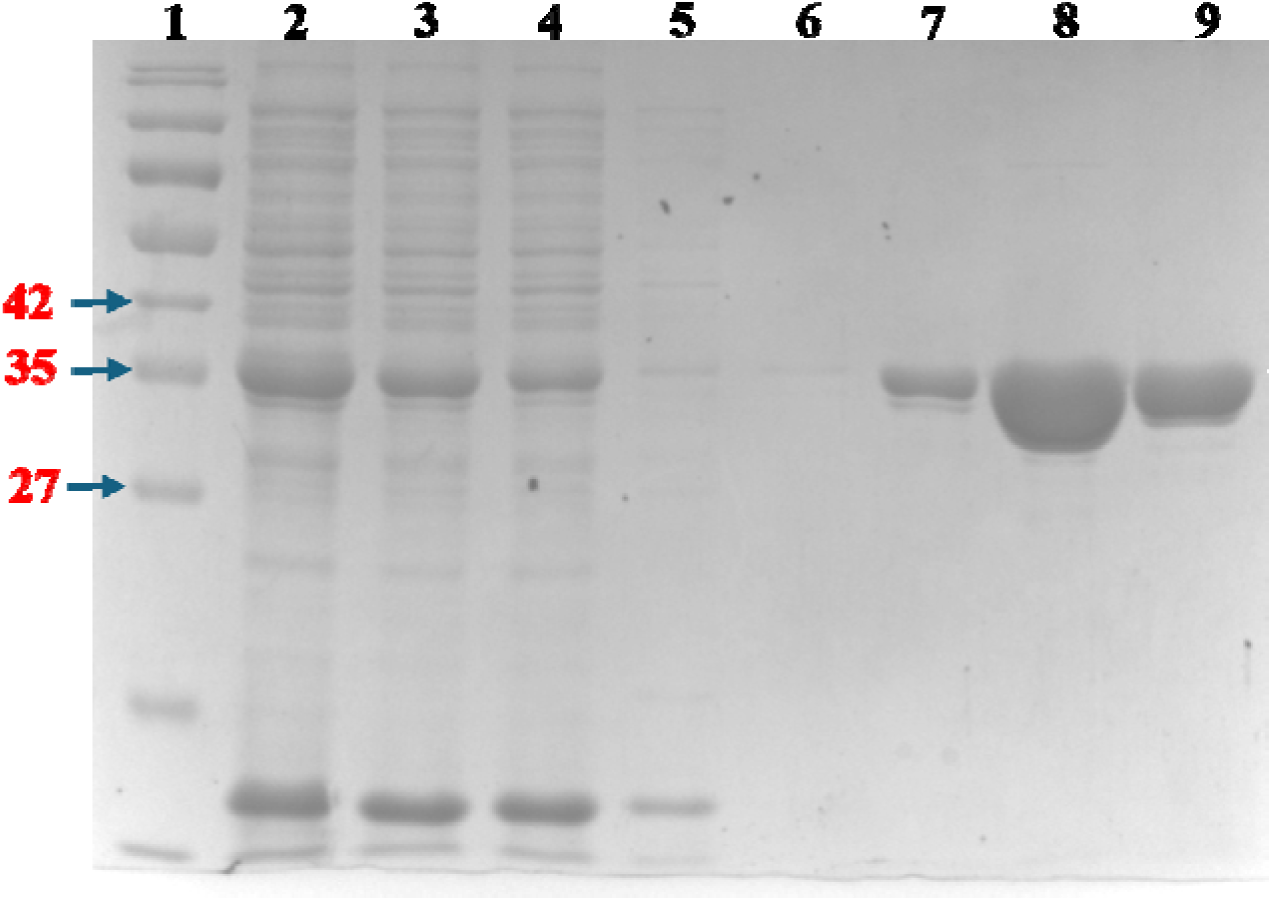
Purification profile of YghA. Numbers on the left indicate the molecular weight (in kDa) corresponding to the respective band. Lane 1 – 180 kDa ladder; Lane 2 – lysate; Lane 3 – supernatant after clarification of cell debris; Lane 4 – flowthrough after column/Ni-NTA binding; Lane 5 – first wash; Lane 6 – second wash; Lane 7 – eluate 1; Lane 8 – eluate 2; Lane 9 – eluate 3.

### 3.2 Sequence and structure analysis of YghA

Although the crystal structure of *E. coli* YghA has not been determined experimentally, the crystal structure of YghA in *Salmonella enterica* in complex with NADH has been solved via X-Ray diffraction at a resolution of 1.25Å (PDB id: 3R3S). Sequence alignment revealed that both proteins are very similar, sharing over 93% amino acid identity (Figure 2A). In agreement with the sequence homology, the structures of *S. enterica* YghA and *E. coli* YghA, as predicted via AlphaFold, are highly similar (Fig. 2B). *S. enterica* YghA crystalizes as a homotetramer (Figure 2C) with each single domain monomer capable of binding one molecule of NADH. Cofactor binding is facilitated by several amino acids forming polar contacts within the binding pocket (Figure 2D). As the sequence and structure of *E. coli* and *S. enterica* YghA are near identical, such properties are likely shared between the two proteins. Further analysis of the *E. coli* YghA amino acid sequence revealed sequence motifs associated with proteins capable of binding to both NADH and NADPH (50). This observation is consistent with previously reported experimental data suggesting that *E. coli* YghA can bind to either cofactor (42).

**Figure 2.**
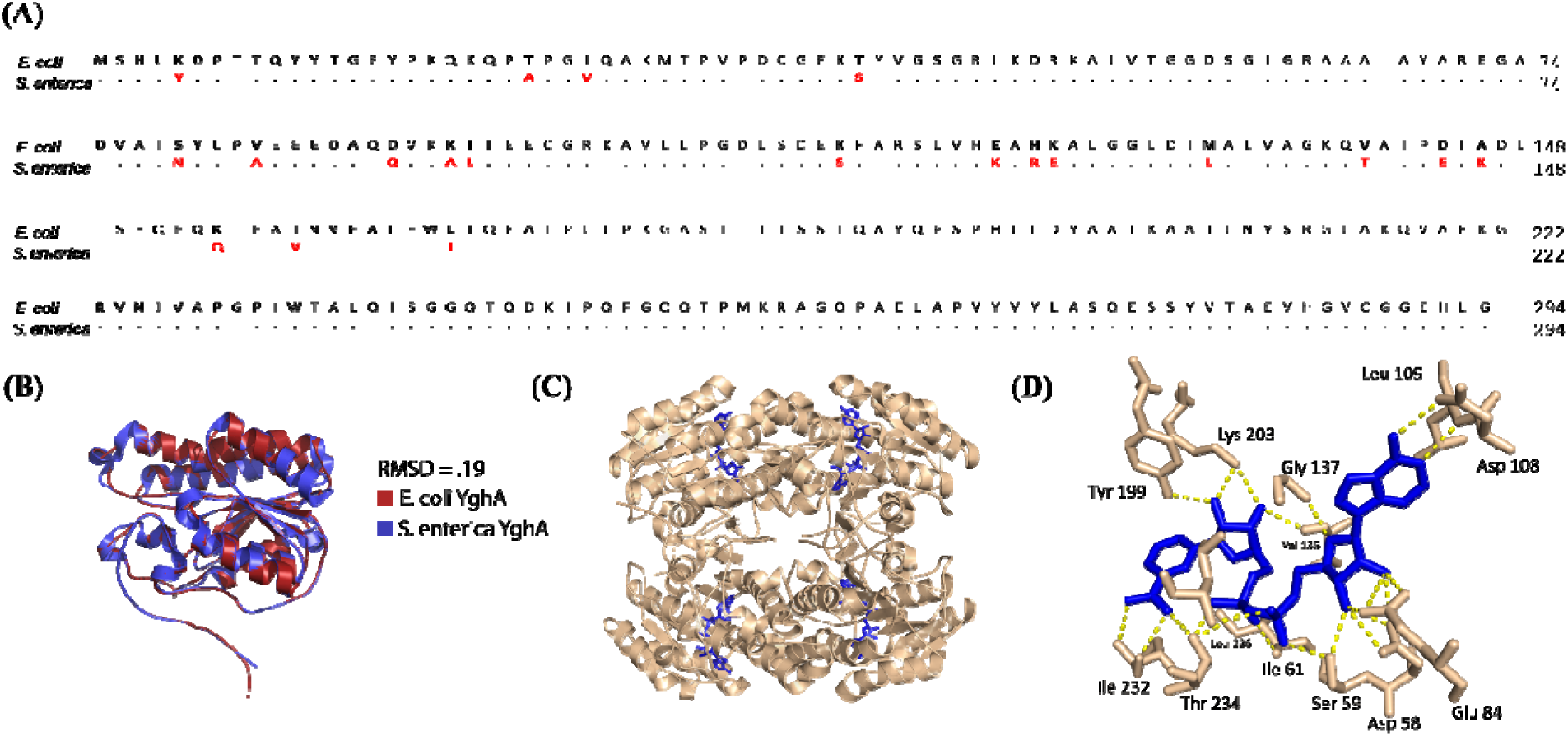
(A) Sequence alignment of E. coli and S. enterica YghA. (B) Overlay of the AlphaFold predicted E. coli Ygha and the X-ray crystal structure of S. enterica YghA. (C) S. enterica YghA tetramer, represented as tan cartoon, bound to NADH, represented as blue licorice. (D) NADH binding pocket of S. enterica YghA. Polar contacts between YghA (tan) and NADH (blue) are represented by yellow dashed lines.

### 3.3 UV-vis absorption spectroscopy

UV-vis absorption is commonly used to study structural changes in the proteins as a function of pH (51,52). The absorption pattern at 278 nm has been ascribed to the π – π* transition of the aromatic amino acid residues Tyr, Trp and Phe for the classical well defined BSA protein (53) and reflects major alpha-helical content (54–56). Amino acid sequence analysis also revealed that the aromatic content of each monomer of *E. coli* YghA possesses 12 tyrosine, 6 phenylalanine and 2 tryptophan residues. The absorption pattern of YghA observed for the wavelengths around 278 nm reflects existence of alpha-helical structure (Figure 3A). YghA appears to be structurally sensitive to pH as changes in the pH of the medium correlated with the variation in UV signal. This suggests that pH induced structural changes expose the aromatic amino acid residues buried in the tertiary structure and alter absorption intensity at 278 nm (57). To investigate further the variation in UV signal we analyzed YghA with relatively more sensitive fluorescence measurements.

**Figure 3.**
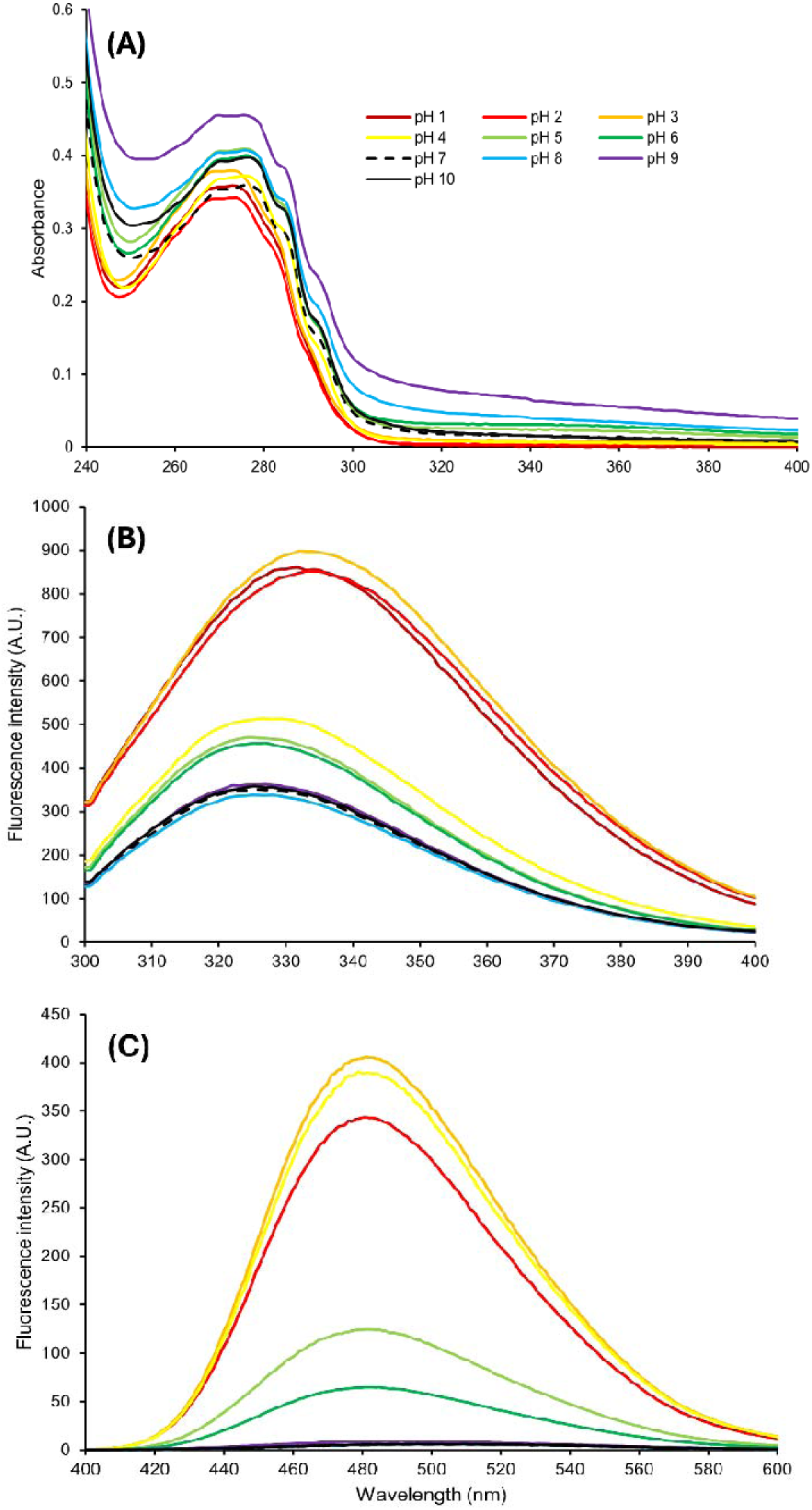
Variation in the optical characteristics of YghA as a function of pH at 298K. The reference pH 7 observation is denoted with a dashed line.(A) UV absorption spectra (B) fluorescence spectra and (C) emission spectra of ANS.

### 3.4 Fluorescence study

Relative to the UV-Vis absorption, fluorescence measurements can reveal greater insights into protein conformation. For example, structural perturbations due to change in the micro-environment of the aromatic rings of tyrosine and tryptophan can shift the emission spectrum towards higher wavelengths (56,58). In agreement with UV-Vis absorption data, YghA displayed pH based variation in fluorescence intensity (Figure 3B). When pH of the buffer was increased to 8,9 and 10, no remarkable changes in the fluorescence intensity as compared to circumneutral pH 7 was observed. However, at acidic pH, we observed an increase in fluorescence intensity with concomitant decrease in pH. A prominent red shift was also observed in acidic pH environment. pH treatments of 6, 5 and 4 yielded similar intensities with pH 4 showing a near 1.5-fold increase as compared to the value observed at the neutral pH. The values increased 2.5-fold when the pH was lowered to values of 3 and below. The fluorescence intensity observed for pH treatment of 3, 2 and 1 were comparable. The data suggests that structural integrity of YghA is relatively resilient to the alkaline pH values while in acidic environment the changes appear to be more abrupt.

### 3.5 ANS study

8-anilino-1-naphthalenesulfonic acid (ANS) is an amphiphilic fluorescent probe which is commonly used to investigate the changes in the protein conformation. The sulfonate group of ANS interacts with the cationic groups of hydrophobic amino acids lysine and arginine and enhances fluorescence intensity along with fluorescence shift towards blue spectrum (59). We used ANS to investigate the change in fluorescence pattern of YghA under different pH treatments (Figure 3C). The increase in intensity and shift to a lower wavelength (blue shift) of fluorescence in presence of the dye was attributed to its interaction with the amino acid residues of the exposed hydrophobic core of the protein. We observed that at circumneutral pH 7, the fluorescence intensity was barely detectable. As compared to pH 7, there was a 68 to 79.4-fold increase in fluorescence intensity in pH 2-4 treatments. While the corresponding increase in the fluorescence intensity at alkaline pH 8-10 treatments was approximately between 1 to 2-fold only. This suggests that at lower pH the ionic bonds are susceptible to disruption and lead to exposure of hydrophobic core of the protein to the hydrophilic ANS probe present in the buffer solution with concomitant increase in the fluorescence intensity of the solution.

### 3.6 Structural analysis of YghA by circular dichroism (CD)

CD has been widely used to access structural stability of proteins (55,60–62). We assessed the secondary structure properties of YghA using CD as a function of pH (Figure 4). At the circumneutral pH, the secondary structure of the protein is primarily composed of alpha-helix with a characteristic maximum absorption observed at 222 nm and a less negative absorption at 208 nm which is due to n to π and π to π transition, respectively (54,63). It is in concurrence with the earlier structural prediction form the amino acid sequences. At pH 8 and 10 an increase in alpha-helical content of YghA was observed with an increase in the negative ellipticity while at pH 9 a decrease of same was observed. Similarly with a decrease in the pH values to 6 and 5 there was an increase in alpha-helical content and at pH value 4 it was comparable to that of pH 7. At pH values 3 and lower there was an abrupt loss of the alpha-helical structure.

**Figure 4.**
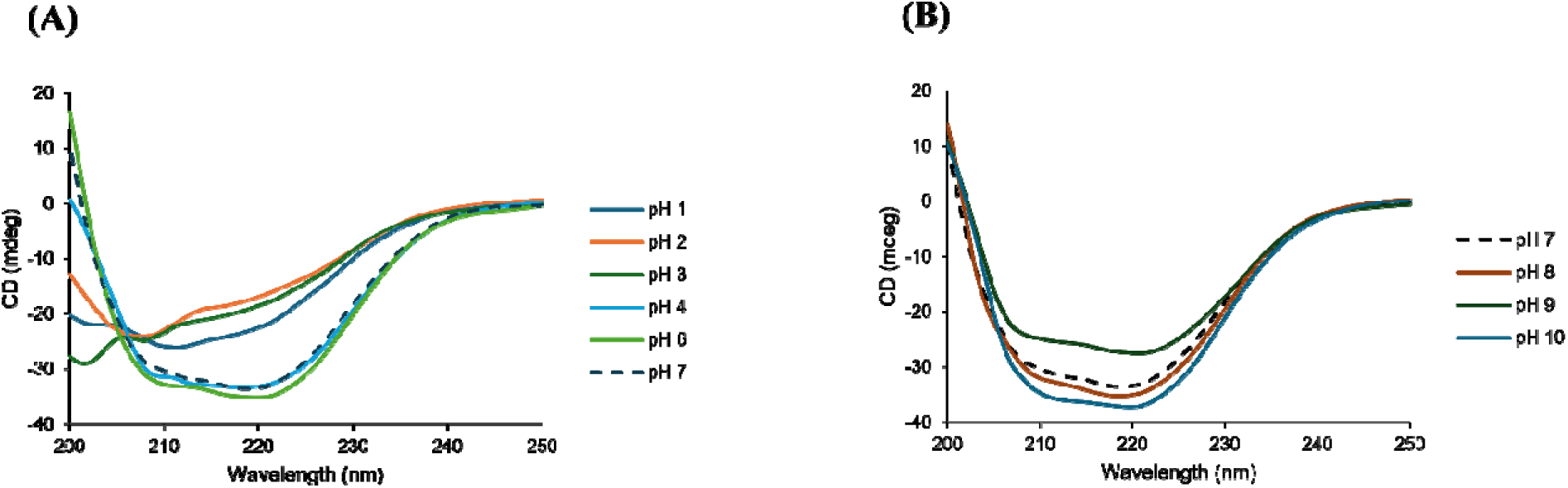
Far-UV CD spectra of YghA under (A) acidic pH conditions (B) basic pH conditions. The reference pH 7 spectra is indicated in dashed line.

### 3.7 Docking and Molecular Dynamics Simulation studies

We computationally investigated the interactions between *E. coli* YghA and each of the cofactors. The Rosie Ligand docking protocol predicts that NADH formed a relatively more stable complex with YghA compared to NADPH (Rosie interface energies: NADH -23.189, NADPH -10.973). FitDock (45) also predicted that NADH is more preferred by YghA (Vina docking energies: -12.5 for NADH and -6.2 for NADPH). The interaction of YghA with respective cofactor is visualized in Supplementary Figure S1.

We simulated to test the influence of pH on interactions between YghA and respective cofactors. The pulling simulation findings indicate that a notably greater force (kJ/mol) and time (ps) were needed to extract NADH from the YghA pocket than NADPH under the studied pH conditions, which indicate that NADH is more preferred. Notably, there is a strong concordance between the docking and pulling simulation results and supports the earlier experimental findings of NADH being the preferred cofactor (Supplementary Figure S2).

Results of simulations also suggest a pH-dependent variations in the force and time required to extract NADH and NADPH from the binding pocket in YghA. It suggests influence of the pH on the YghA-cofactor interaction. In contrast to stronger interactions at pH 7, lower force interaction at pH 1 indicates weaker and unstable binding for NADH, while for NADPH the weaker force exerted was at pH 10 (TABLE 1).

**Table 1:** Pulling forces exerted on NADH and NADPH at different pH levels.

| pH | NADH |  | NADPH |  |
| --- | --- | --- | --- | --- |
|  | Time (ps) | Force (kJ/mol/nm) | Time (ps) | Force (kJ/mol/nm) |
| 1 | 300 | 2750 | 300 | 2500 |
| 4 | 490 | 4500 | 300 | 2800 |
| 7 | 510 | 4800 | 310 | 2900 |
| 10 | 415 | 3800 | 270 | 2100 |
| 12 | 395 | 3500 | 300 | 2650 |

We also tested the structural stability of YghA under different pH treatments (Figure 5). Consistent and equilibrated variations in the backbone root mean square deviation (RMSD) values were noted at neutral and basic pH 12, respectively, during the 100 nanosecond (ns) MD simulations. Under tested conditions, YghA tetramer preserves consistent and stable structural properties throughout the simulation time. The tetramer structure is well-equilibrated and does not significantly deviate from its initial shape. While significant variations in the RMSD of the YghA tetramer backbone were noted at pH 1 (Figure 5A). This suggests that the stability and conformational dynamics of the tetramer structure are specifically impacted by the acidic environment. The variations could be explained by the higher structural flexibility and conformational rearrangements brought about by the protonation of amino acid residues, or modifications of the hydrogen bonding patterns within the protein.

**Figure 5.**
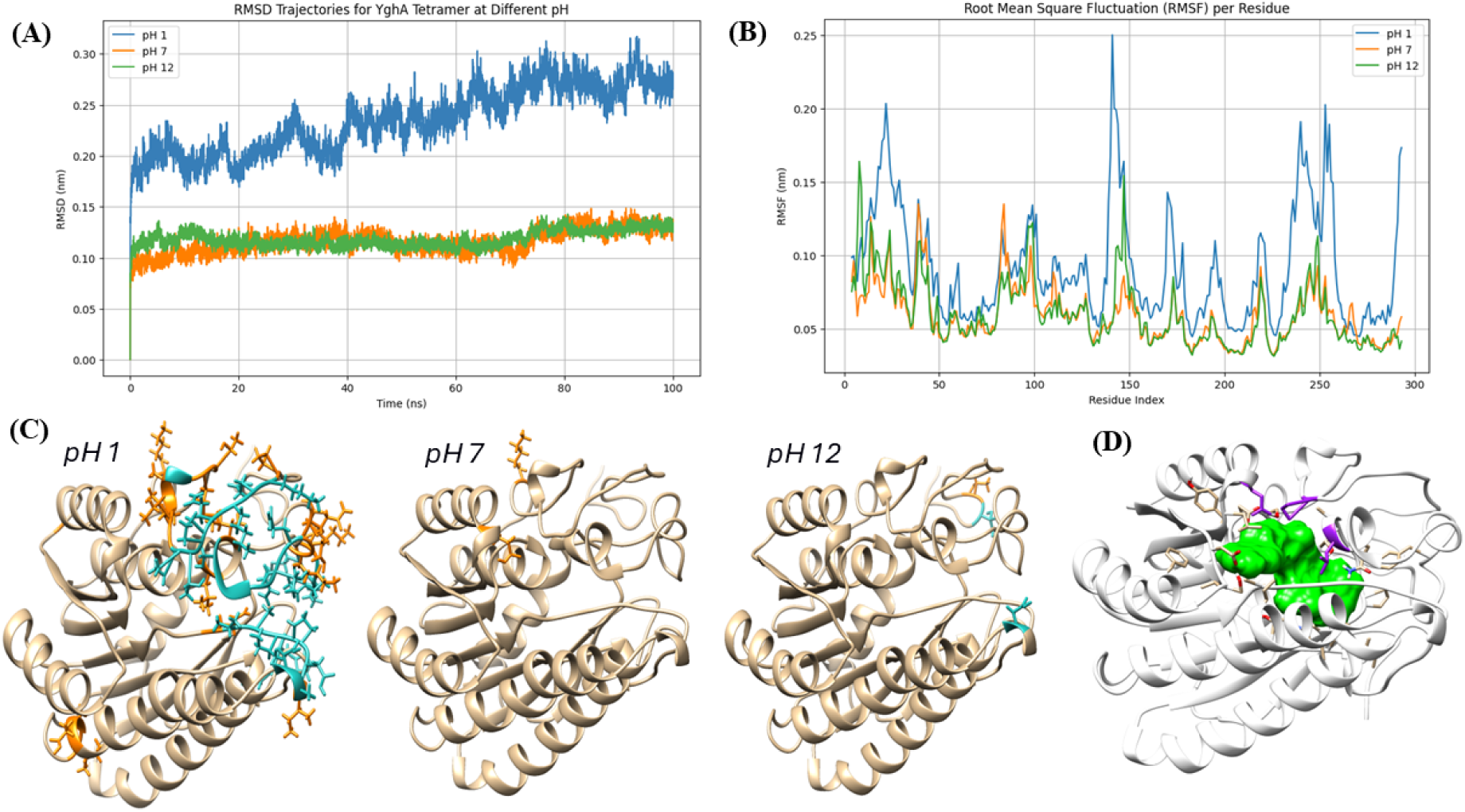
Stability of E. coli YghA at different pH treatments. (A) RMSD of YghA tetramer backbone during MD simulations (B) Root Mean Square Fluctuation (RMSF) analysis per residue of YghA dynamics at different pH conditions. (C) Flexible amino acids during 100ns of simulations aa different pH levels. Blue shows more flexibility than the orange ones. (D) Flexible pocket residue in purple, PRO 24 and LEU 236 at pH:1 and GLH 84 (protonated GLU) at pH:7. Green surface shows the binding cavity of YghA for cofactors.

The flexibility and stability of amino acid residues offer insights into the possible structural modifications to the protein. Under tested pH conditions, the Root Mean Square Fluctuation (RMSF) analysis per residue shows differences in residue flexibility (Figure 5B). At extreme pH values, some residues show enhanced flexibility and most likely contribute to the loss of structural thermodynamic homeostasis. This increased fluidity around some amino acid residues could explain the CD signal corresponding to the loss of the alpha-helix structure in Figure 4. GLH 84 (protonated GLU) at pH 7 and PRO 24 and LEU 236 at pH 1 are two examples of the flexible pocket residues that are important for YghA’s binding cavity (Figure 5C, D). These residues show pH-dependent flexibility, which may have an impact on the protein’s ability to bind and maintain its structural integrity.

Results of simulation studies demonstrate the sensitivity of the YghA tetramer to pH variations with significant deviations in the backbone RMSD and RMSF values under acidic conditions as compared to the neutral and basic pH environments.

## 4. Conclusions

In this study, we report a previously uncharacterized protein which has relevance in bioenergy studies. YghA protein from *E. coli* displays both sequence and structural similarity to its counterpart reported from *Salmonella enterica*. The structural integrity of YghA is relatively resilient to an increase in pH as compared to acidic pH and this inference is drawn from multiple observations in this study. As compared to fluorescence at neutral pH, the fluorescence intensity of YghA increases 2.5-fold at pH 3 and lower. Significant interaction of the ANS dye with the hydrophobic core was observed at pH 2-4 where almost 68 to 79.4-fold increase in fluorescence intensity was observed as compared to pH 7. While at basic pH the increase in signal intensity was only 1-2 fold. Results of the CD studies suggest that YghA is primarily composed of alpha-helix and the secondary structure is abruptly lost when the pH is lowered to 3 and below. Results of docking studies suggested that YghA forms a more stable complex with NADH as compared to NADPH. At neutral pH, YghA displayed higher pulling force in presence of NADH as compared to NADPH. The strength of interaction with cofactors as evident by the pulling force also varied as a function of pH. At pH 1, the low pulling force of NADH as compared to pH 7 suggested a weak interaction while at pH 10, the NADPH exerted a weaker force. Simulation studies also suggested that the structural integrity of YghA is maintained at neutral and basic pH values while at acidic pH significant variations in RMSD value suggest that the conformational stability of tetramer is compromised. The RMSF analysis suggested that GLH 84 at pH 7; PRO 24 and LEU 236 at pH 1 are the flexible residues in the YghA binding cavity and potentially influence the catalytic activity. These residues are promising target(s) for mutagenesis to explore a relatively more stable structure of YghA tetramer which is resilient to acidic pH conditions.

YghA protein holds significant potential to be utilized as a candidate for enzyme immobilization studies for detoxification of furfural from the low pH lignocellulosic hydrolysate before it is utilized as a medium for microbial cultivation in an industrial bioreactor. Since *E. coli* is considered to be GRAS organism, any biomolecule derived from this microbe would not face significant regulatory hurdles in its application for industrial purposes. However, in the absence of any studies related to the variants of YghA derived from *E. coli*, it is difficult to evaluate the results of this study from industrial perspective. It is desirable to pursue further mutagenesis studies targeted towards increasing the tolerance of YghA towards acidic pH. In published studies removal of furfural from lignocellulosic hydrolysate was achieved by in vivo overexpression of FucO and YghA. Acid tolerant version of YghA protein will require evaluation of its activity under hydrolysate conditions in presence of cofactor NADH required for its activity. For continuous catalytic conversion by YghA a steady supply of NADH cofactor will be required which can potentially drive up the costs of hydrolysate detoxification under industrial settings. One potential remedy could be to use a cofactor regeneration system with a NAD^+^ reducing enzyme like formate dehydrogenase (Fdh) from *Candida boidinii*. Fdh can be co-immobilized alongwith acid tolerant version of YghA enzyme to potentially reduce the cost of continuous NADH supplementation. Fdh from *C. boidinii* is resilient to pH variations and has been reported to be used in cell free system for regeneration of NADH with high efficiency (64).

## Acknowledgements

YJC is supported by the National Research Foundation of Korea (NRF), funded by the Korean government (MSIT) (NRF-2020R1A5A2031185, NRF-2020M3A9G3080281 and NRF-2021R1F1A1063769). NIAID, NIH and Department of Health and Human Services are kindly acknowledged for supporting Center for Structural Genomics and Infectious Disease (CSGID) under contract number HHSN272200700058C, HHSN272201200026C and HHSN272201700060C. CSGID is kindly acknowledged for determining the crystal structure of YghA in *Salmonella enterica*.

---

**Figure S1.**
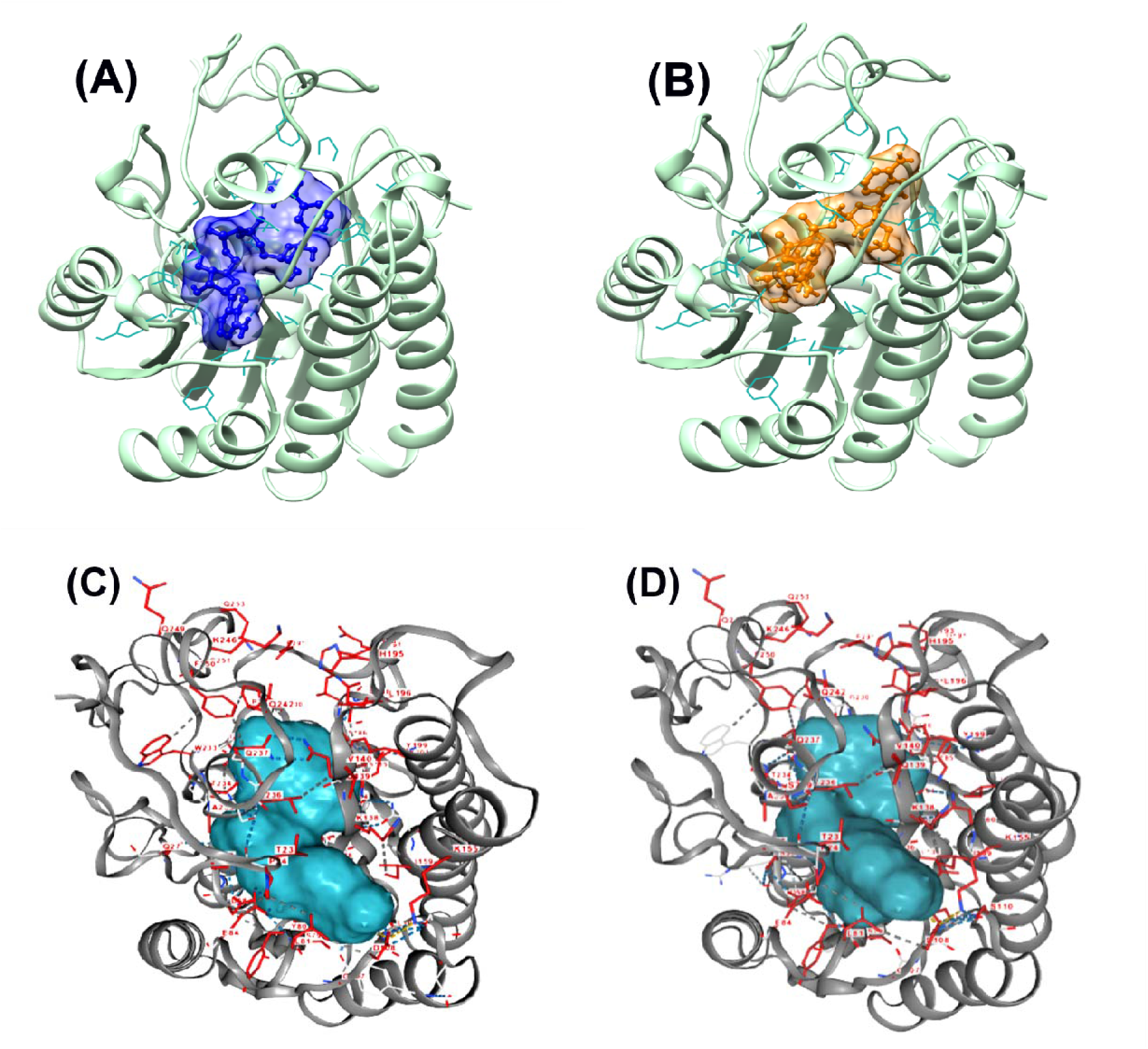
Best docked conformations. (A) NADH in the binding pocket with -23.189 interface energy (B) NADPH in the binding pocket of YghA monomer with -10.973 interface energy. Best docked conformations from FitDock (C) NADH in the binding pocket with -12.5 energy (D) NADPH in the same binding pocket of YghA monomer with -6.2 energy.

**Figure S2:**
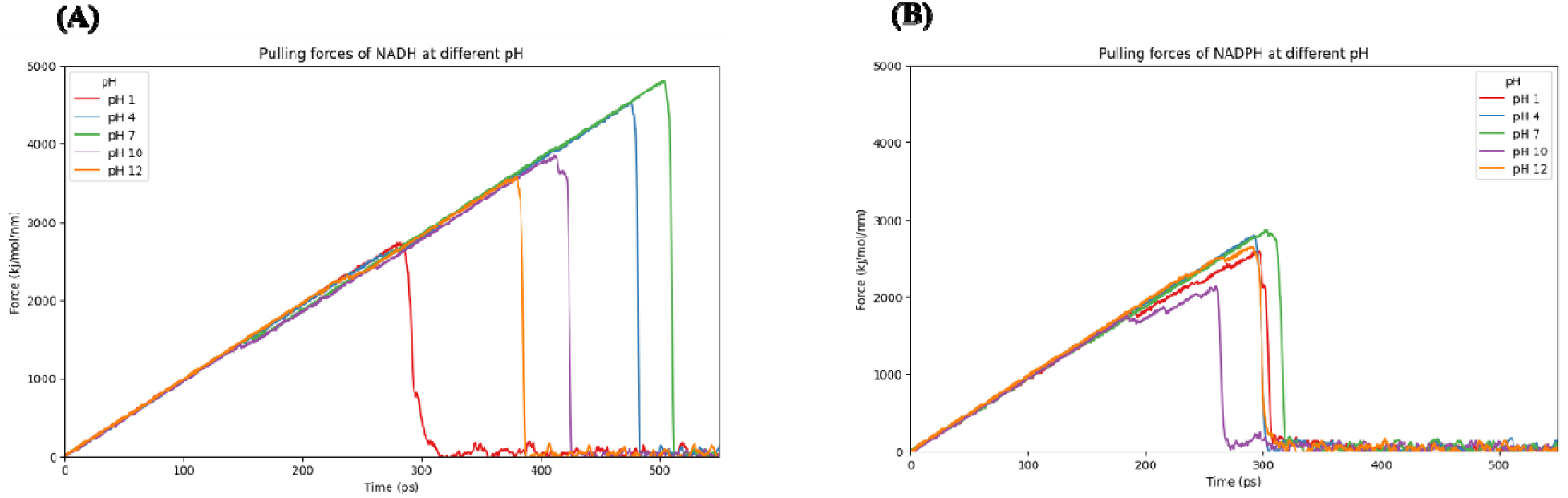
Pull forces exerted on NADH at different pH levels. Pull forces exerted on NADPH at different pH levels.

