## Supplementary data for "Changes in biophysical characteristics of YghA from *E. coli* due to variation in pH"

**Figure S1.** Best docked conformations. (A) NADH in the binding pocket with -23.189 interface energy (B) NADPH in the binding pocket of YghA monomer with -10.973 interface energy. Best docked conformations from FitDock (C) NADH in the binding pocket with -12.5 energy (D) NADPH in the same binding pocket of YghA monomer with -6.2 energy.

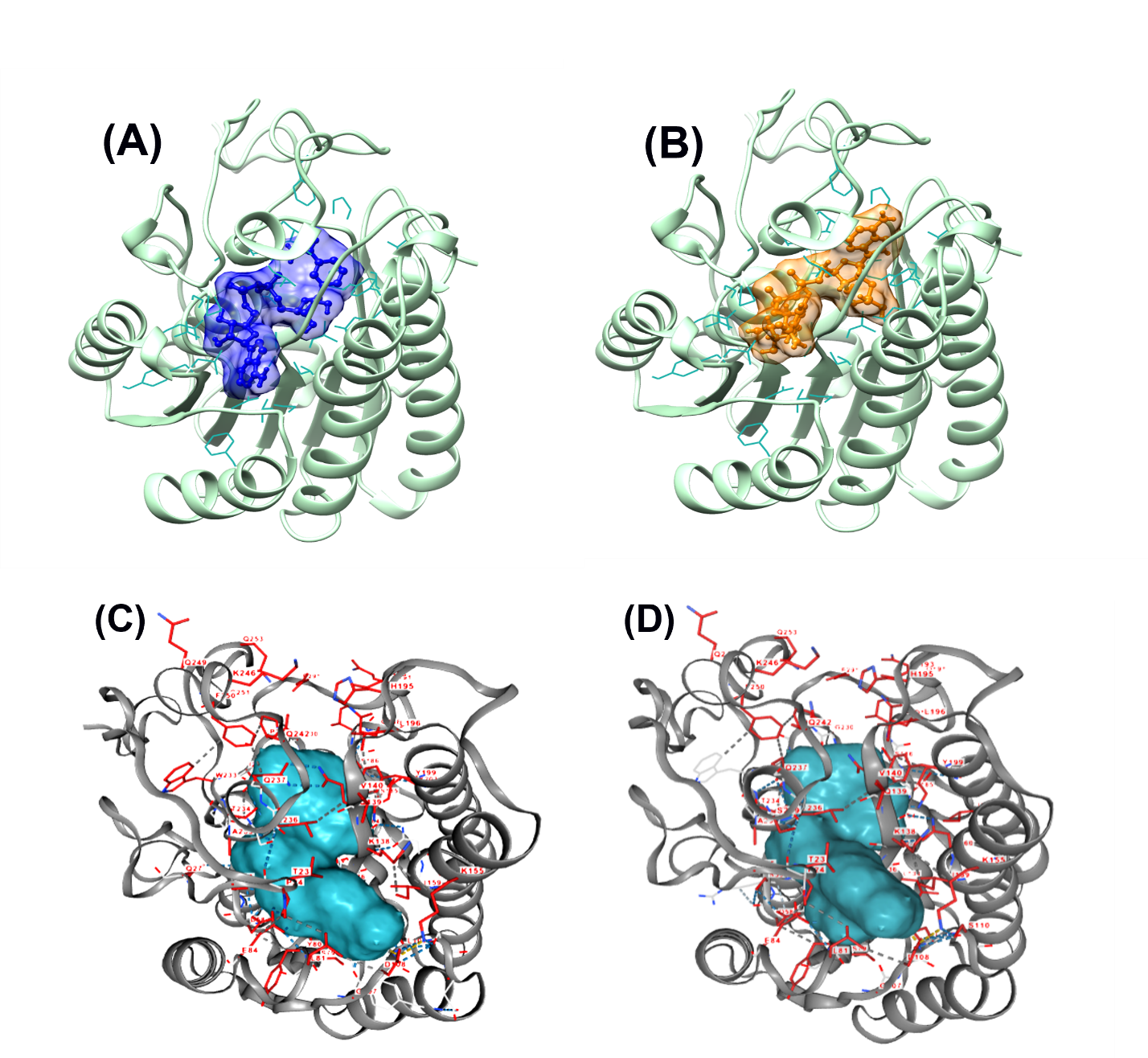

**Figure S2:** Pull forces exerted on NADH at different pH levels. Pull forces exerted on NADPH at different pH levels.

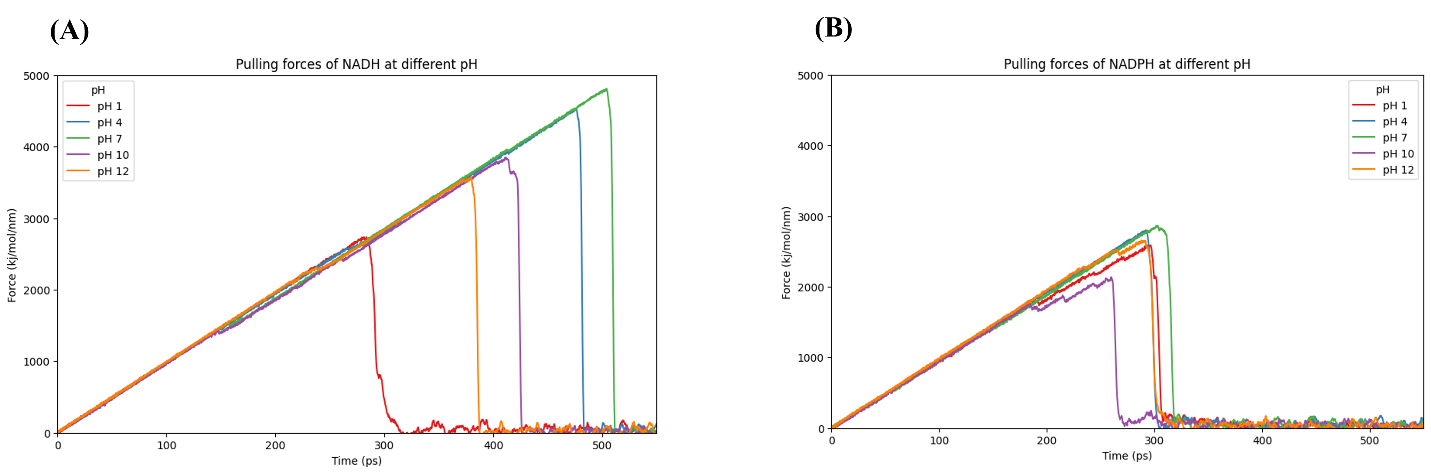
